# The Drosophila genes *disconnected* and *disco-related* together specify development of adult legs

**DOI:** 10.1101/052811

**Authors:** Juan B. Rosario, James W. Mahaffey

**Author notes:** Current address: Integra Labs, Durham North Carolina, United States of America (Juan B. Rosario).

## Abstract

In the fruit fly, *Drosophila melanogaster*, specification of the legs begins during embryogenesis when Wingless signaling induces small groups of cells to form the imaginal disc primordia in the thoracic segments. This signal initiates expression of transcription factors that will later be used to pattern the legs. The paralogous genes *disconnected* and *disco-related* encode transcription factors that are expressed in the disc primordia during early embryogenesis, and their expression continues in the leg discs during larval and pupal stages. The importance of these two genes in establishing the leg development trajectory was indicated by our previous observation that ectopic expression of either gene in the wing discs cells caused legs to develop in place of wings. However, because of their redundancy and requirement for survival during embryogenesis, we were unable to define their role in development of the adult legs. Here, we report loss-of-function analyses of the *disco* genes during development of the legs. We discovered that loss of both genes’ functions causes both truncation of the distal leg with apparent overgrowth of proximal regions and complete loss of legs and ventral thoracic body patterning. At the molecular level we noted reduction or loss of signaling and transcription factors that pattern the proximal-distal axis of the legs. We conclude from these studies that the *disco* genes promote leg development through regulation of signaling processes, but also by stabilizing expression of the leg determination gene network.

## INTRODUCTION

In holometabolous insects, those that undergo complete metamorphosis such as *Drosophila melanogaster*, most adult body structures develop from imaginal discs, groups of cells that are set-aside during embryogenesis, proliferate during larval stages, and form the appendages (mouthparts, legs, wings, halteres, antennae, etc.) and body wall of the adult insect during the pupal stage. The thoracic imaginal discs form during embryogenesis when segmental expression of the morphogen Wingless (Wg) induces a small group of cells to activate appendage-specific gene networks (Cohen et al., 1993; Cohen, 1990; Estella and Mann, 2010; Estella et al., 2003). The dorsal limit of the disc primordia is set through repression by Decapentaplegic (Dpp) and the ventral limit by repression from the Epidermal Growth Factor Receptor (EGFR) (Cohen et al., 1993; Goto and Hayashi, 1997; Kubota et al., 2000; Raz and Shilo, 1993).

Shortly after induction, the thoracic disc primordia separate into the dorsal (wings and halteres) and ventral (legs) imaginal precursors (Cohen et al., 1993). As the dorsal and ventral disc primordia separate, changes in gene expression occur that begin the process of patterning the adult legs; the current model is summarized in Fig. 1. Cells in the posterior compartment of the leg discs express *engrailed* (*en*), which activates *hedgehog* (*hh*) signaling. Cells just anterior to the posterior compartment border receive this Hh signal and respond by activating either *dpp* if dorsal or *wg* if ventral (Basler and Struhl, 1994; Brook and Cohen, 1996; Brook et al., 1996; Jiang and Struhl, 1996; Lecuit and Cohen, 1997; Mohler, 1988; Tabata et al., 1992; Tabata and Kornberg, 1994; Theisen et al., 1996). In this manner the overall axes of the discs and of the adult legs are determined. Cells in the center of the disc perceive high collective concentrations of Wg and Dpp and initiate expression of *Distal-less* (*Dll*) (Fig. 1B). Although *Dll* is expressed throughout the thoracic disc primordia during initiation in the early embryo, this later *Dll* expression is restricted to cells that will form the medial-to-distal portion of the legs (Angelini and Kaufman, 2005; Bolinger and Boekhoff-Falk, 2005; McKay et al., 2009; Panganiban, 2000b). Cells in the outer region of the discs do not respond to the Hh, Wg and Dpp signals, and these cells activate genes necessary for proximal identities, genes such as *homothorax* (*hth*) and *teashirt* (*tsh*) (Diaz-Benjumea et al., 1994; Estella et al., 2012; Gonzalez-Crespo et al., 1998; Gonzalez-Crespo and Morata, 1996). Cells that lie between these two domains express *dachshund* (*dac*), which defines the medial region (Giorgianni and Mann, 2011; Mardon et al., 1994). In this manner three major proximal to distal domains of the adult leg are established in the larval imaginal discs. The Hth/Tsh domain determines the body wall and the most proximal segment of the leg (the coxa); Dll specifies the distal portion of the leg, and Dac the medial.

**Figure 1:**
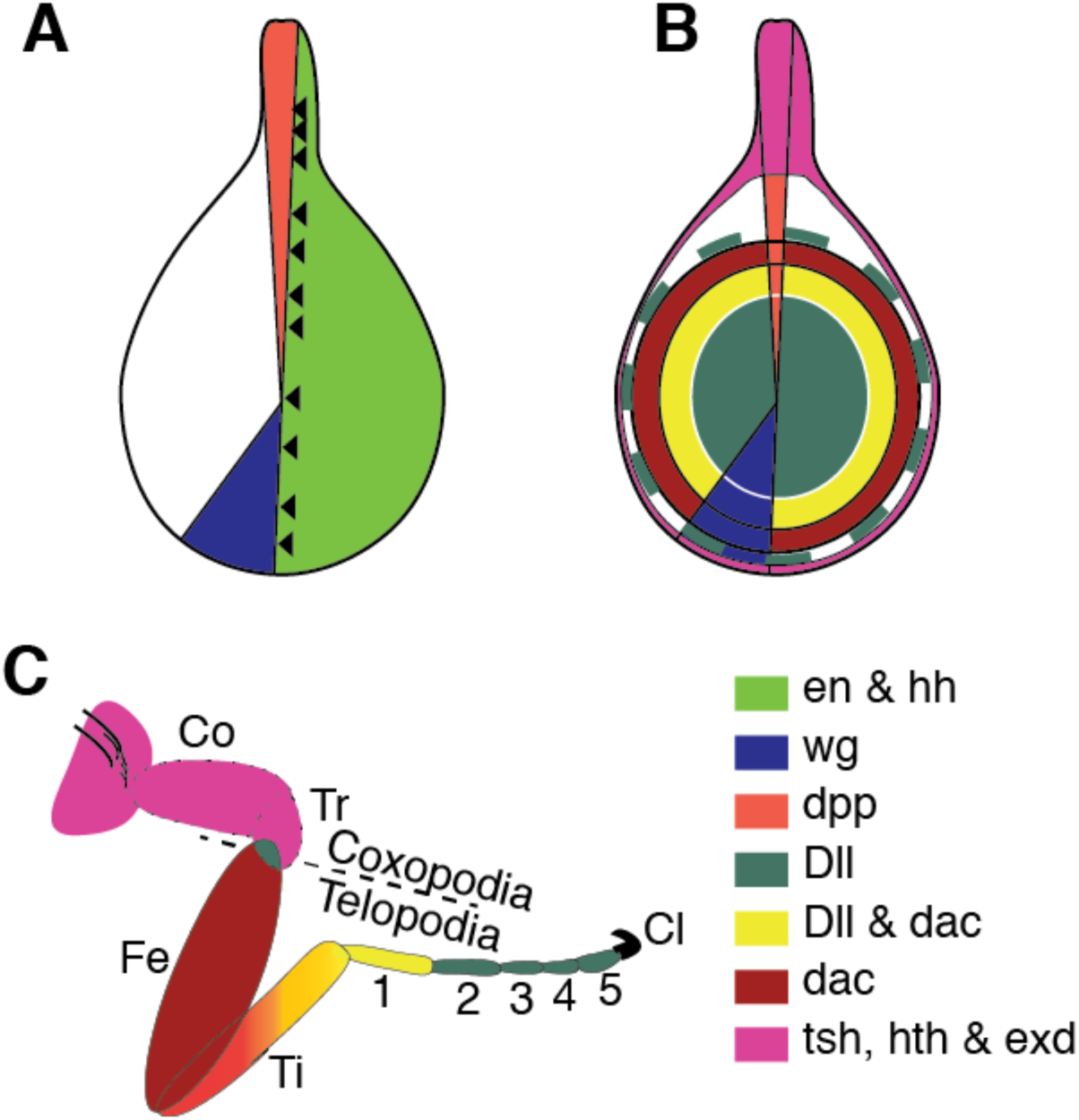
Model of genetic specification of Drosophila leg development (adapted from that described by Lecuit and Cohen, 1997 and Giorgianni and Mann, 2011. (A) The segment polarity gene *engrailed* (*en*) is expressed in the cells of the posterior compartment of the leg disc and activates *hedgehog* (*hh*, bright green). Cells anterior to the compartment boundary receiving Hh signal (arrowheads) activate *decapentaplegic* (*dpp*) dorsally (orange) and *wingless* (*wg*) ventrally (blue). (B) In the center of the disc *Distal-less* (*Dll*) (dark green) is activated in response to high combined concentration of Wg and Dpp. Cells away from that center express *dachshund* (*dac*) and *Dll* (yellow) while further cells express only *dac* (red). Cells producing the most proximal structures and the body wall express teashirt (*tsh*) and homothorax (*hth*) (magenta). (C) Link between the adult structure and the corresponding genetic domains in the imaginal disc.

Other genes are expressed in the imaginal discs primordia during early embryogenesis, such as the paralogous pair *disconnected* (*disco*) and *disco-related* (*disco-r*) (Cohen et al., 1991; Lee et al., 1991; Mahaffey et al., 2001), together referred to as “the *disco* genes” hereafter. The *disco* genes encode redundant C2H2 zinc finger transcription factors (Mahaffey et al., 2001). They are initially expressed in the thoracic disc primordia during embryonic stage 11 (Cohen et al., 1991; Lee et al., 1991; Mahaffey et al., 2001), and they are continually expressed in the ventral imaginal discs after separation of the dorsal discs precursors. By late third instar the *disco* genes are expressed throughout most of the cells of the leg imaginal discs, encompassing the *Dll* and *dac* domains and extending into more proximal regions (Patel et al., 2007). The importance of the *disco* genes during leg specification was demonstrated by ectopic expression experiments. Expression of either gene in the wing discs causes these discs to form legs (Patel et al., 2007). However, what role these genes have during normal leg development remained elusive, since loss of both genes is lethal to embryos, and randomly generated clones of cells lacking both genes are rarely recovered in the leg discs (Patel et al., 2007). Below we describe results of our investigation into the role of the *disco* genes during development of the Drosophila legs. We conclude that these genes are necessary to stabilize leg development, likely through stabilization of the leg transcriptional network.

## MATERIALS AND METHODS

### Fly stocks

Oregon-R was used as a wild type. The following fly lines were obtained form the Bloomington Stock Center: *Df (1) ED7355* (FBst0008899, Ryder et al., 2004); *tubP-GAL80*^*ts*^ (FBst0007108, McGuire et al., 2003); the *disco-r RNAi line y[1] sc[*] v[1]; P[y[+t7.7] v[+t1.8]=TRiP.HMS02247]attP2* from the Transgenic RNAi Project (FBst0041683); *Act5C-Gal4* (FBst0004414); *Dll*^*md23*^ (FBst0003038, Calleja et al., 1996). We used several lines carrying the *disco*^*1*^ allele (Fischbach and Heisenberg, 1984), one from A. Campos (McMaster University), one from J. Hall (Brandeis University) and one from the Bloomington Drosophila Stock Center (FBst0005682). Three additional *disco-r* RNAi lines were obtained from the Vienna Drosophila RNAi Center (VDRC), *P[KK103892]VIE-260B* (FBst0473452), *w[1118]; P[GD13583]v35750* (FBst0461319), *w[1118]; P[GD13583]v35751* (FBst0461320). *UAS-GFP/UAS-GFP* was a gift from Dr. Patricia Estes, North Carolina State University. Flies were raised on standard cornmeal-agar-molasses media at 17°C except where otherwise required by the experiment.

### Fly crosses to generate *disco*-lof

To generate *disco*^*1*^/*disco*^*1*^; *disco-r* RNAi/*disco-r* RNAi flies, we crossed females homozygous for *disco*^*1*^ to males *Sco/ CyO, Act-GFP*. We selected male *disco*^*1*^/Y; +*/ CyO, Act-GFP* and crossed these to *disco*^*1*^ homozygous females. We selected females *disco*^*1*^/*disco*^*1*^; *CyO, Act-GFP*/+ and crossed them to males homozygous for *disco-r* RNAi line, *P[KK103892]VIE-260B*. From this cross we selected males *disco*^*1*^/Y; *disco-r* RNAi /*CyO, Act-GFP* and crossed these to *disco*^*1*^/*disco*^*1*^; *CyO, Act-GFP*/+ females. Pair matings of the *CyO, Act-GFP* progeny were set up, and once larval activity was noted in the vial, DNA was extracted from the parents and amplified to verify the presence of the RNAi construct in both parents using the primer sequences recommended by the VDRC. To activate the *disco-r* RNAi, *disco*^*1*^/*disco*^*1*^; *disco-r* RNAi/*disco-r* RNAi females were crossed to +/Y; *Act-Gal4/ CyO, Act-GFP*. Non-GFP male larvae were selected by the presence of the testis. In some cases we used *tub-Gal80*^*ts*^ to repress the Gal4 system until larval stages (McGuire et al., 2003).

### Testing other RNAi lines and an additional *Gal4* driver

For *Dll-Gal4* activation of the RNAi, we generated a *Dll-Gal4*, *tub-Gal80* fly line by recombination of *P[w[+mW.hs]=GawB]Dll[md23]* with *w[*]; P[w[+mC]=tubP-GAL80[ts]]10*. These flies were crossed to homozygous *disco*^*1*^ females to generate *disco*^*1*^/Y; *Dll-Gal4, tub-Gal80*^*ts*^/+ flies, which were crossed to the *disco*^*1*^/*disco*^*1*^; *disco-r* RNAi/*disco-r* RNAi. To test different *disco-r* RNAi constructs, *disco*^*1*^/+; *Dll-Gal4, tub-Gal80*^*ts*^/+ females were crossed with males homozygous for the *disco-r* RNAi and male adults or pharate adults were scored. We tested two additional *disco-r* RNAi lines from VDRC, GD35750 and GD35751, and a *disco-r* RNAi line from the Transgenic RNAi Project (TRiP).

### Growth conditions for *disco*-lof

Eggs were collected for two days at 17°C and then the adults removed. Four days later the progeny were transferred to 30°C to inactivate the *tub-Gal80*^*ts*^. Pharate adults (in this case males) were dissected from the pupal case and along with the eclosing males, were stored in 70% ethanol. For molecular analysis we followed a similar time regimen, dissecting discs from late third instar male larvae. RNAi and sibling wild type discs were collected together for *in situ* and immunostaining. We left a portion of the gut attached to the control carcasses to distinguishing them from the *disco*-lof. Note, *tub-Gal80*^*ts*^ was not required to pass embryogenesis with *Act-Gal4*; however, we followed the same timing for elevating the temperature with the *Act-Gal4* and *Dll-Gal4* larvae. Others and we have noted a slight affect on leg development with the *Dll-Gal4* driver since the *Gal4* insertion compromises the *Dll* gene (Calleja et al., 1996; Grubbs et al., 2013). This was taken into account in all analyses.

### Determination of *disco*^*1*^ viability

To determine the viability of *disco*^*1*^, we collected males and females of three different *disco*^*1*^ lines (see Fly stocks, above) and allowed them to mate for 48 hours. We transferred the parents to a collection cup with a grape agar plate for 18 hours. We collected 200 embryos from these plates, and transferred them to fresh grape agar plates and allowed 28 hours for the larvae to hatch. Un-hatched embryos were cleared for cuticle examination to determine the number of unfertilized or undeveloped eggs. The remaining larvae were transferred to standard food diet and allowed to develop. We counted pupae and adults to determine numbers surviving to each stage. The disco genes were sequenced from all lines to verify they carried the *disco*^*1*^ allele.

### Molecular analyses

Embryos were collected on grape plates, and larvae for disc dissections were grown on standard Drosophila medium. Fixation of embryos and discs essentially followed that of (Tautz and Pfeifle, 1989). Embryo and larval cuticle analyses followed that described in (Pederson et al., 1996). The single and double fluorescence *in situs* followed the protocol of (Juarez et al., 2011; Kosman et al., 2004). Antibody detection and double *in situ*/antibody detection is described in (Heffer et al., 2013). Antibodies used, rabbit anti-Tsh (S. Cohen, IMCB, Singapore); anti-Dachshund, mAbdac2-3 (Mardon et al., 1994); anti-Wingless 4D4 (Brook and Cohen, 1996); anti-Beta-gal (Promega); anti-pmad, Phospho-Smad1/5 (Cell Signaling Technology). *Dll*, *disco* and *wg* probes for *in situ* hybridization were described in Mahaffey et al., 2001). TUNEL analysis was performed using ApopTag^®^ Red In Situ Apoptosis Detection Kit (Millipore) following procedure for whole tissue analysis, but scaled down for Drosophila. Fluorescent stained discs were mounted in 70% Glycerol, enzymatic detections in Crystal Mount^®^. Images were acquired using a Zeiss LSM 710 confocal microscope system or Zeiss Axioplan microscope with a Q Imaging Micropublisher 5.0 RTV camera. We used ImageJ for quantification of fluorescence (Schneider et al., 2012). Confocal z-stacks were compressed by SUM, and measurements made from three of the brightest regions, standardized by area. p values obtained using JMP version 10 (SAS Institute Inc., Cary NC, USA). For presentations graphics, Adobe Photoshop and ImageJ were used for brightness/contrast adjustments and cropping. All images in each figure were processed identically. Adobe Illustrator was used to assemble the final figures.

## RESULTS

### Loss of function phenotypes

To assess the role of the *disco* genes during development of the adult legs, we eliminated the function of both genes after embryogenesis by combining a null mutation in *disco* (*disco*^*1*^) (Heilig et al., 1991) with UAS-driven *disco-r* RNAi, controlling spatial and temporal activity of the RNAi during development. We refer to this combination as “*disco*-lof” below. We used two different *Gal4* drivers and four different RNAi constructs with comparable results (see Materials and Methods). Note that homo-or hemizygosity for *disco*^*1*^ was initially reported to be viable (Steller et al., 1987), but there have been reports that it is lethal (Dey et al., 2009; Glossop and Shepherd, 1998). Therefore, we tested three *disco*^*1*^ lines (see Table 1), and found that only the line from the Bloomington Drosophila Stock Center was appreciably less viable than wild type flies. In the present work we used the line we refer to as *disco*^*1*^(C).

**Table 1:** Analysis of *disco*^*1*^ viability (from the total number of hatched embryos) comparing three different *disco*^*1*^ lines.

|  | Ore R | Bloomington | (C) | *w, f* |
| --- | --- | --- | --- | --- |
| Collected | 200 | 200 | 200 | 200 |
| Unfertilized/undeveloped | 19 | 33 | 7 | 14 |
| Total number of developed eggs | 181 | 167 | 193 | 186 |
| Dead embryos | 0 | 20 | 7 | 5 |
| Dead Larvae | 6 | 22 | 20 | 12 |
| Dead pupae | 2 | 33 | 13 | 11 |
| Eclosed<br>(% of total # fertilized eggs) | 173<br>(95.6%) | 92<br>(55.1%) | 153<br>(79.3%) | 158<br>(84.9%) |

We first examined viability and developmental trajectory of the *disco*-lof progeny from the crosses. We shifted larvae at various times to the non-permissive temperature for *tub-Gal80*^*ts*^ (30°C), thus activating the *disco-r* RNAi. Shifting during first or early second instar resulted in larval death, while shifting during third instar resulted in pharate adults with only minor leg defects. Shifting to 30°C during second instar (96-144 hrs at 17°C) generated pharate adults with moderate and severe phenotypes, so this timeframe was used for most experiments.

The normal *Drosophila* leg is composed of five segments (Fig 1C). The body wall and coxa together form the coxapodia, while the trochanter, femur, tibia and tarsus make up the telopodia (Gonzalez-Crespo and Morata, 1996; McKay et al., 2009). The tarsus is further divided into five sub-segments with the pulvilli and claws attached to the most distal of these (t-5). Each leg segment has a characteristic arrangement of bristles, in general organized into longitudinal rows around the circumference of the leg. The bristles have an overall polarity such that most are oriented toward the distal portion of the leg (Hannah-Alava, 1958; Held, 2002; Schubiger et al., 2012; Tokunaga, 1962). On the more distal leg segments, bristles are usually associated with bracts, small, thick trichomes induced by the neighboring bristle. Bristles in the proximal leg segments (up until proximal to mid-femur) are not associated with bracts.

Upon dissecting the *disco*-lof pharate adults from the pupal cases, we noted two main classes of leg defects. Many pharate adults were missing entire legs, while those legs that formed were abnormal (Fig 2B-D). For those legs that were present, *Dll-Gal4*-driven RNAi usually lacked all tarsal segments and had an enlarged fusion of the proximal segments that appeared to be composed of coxa, trochanter and femur (Fig. 2B,C and Fig. 3B,C). Legs from the *Act-Gal4* driver crosses had similar proximal fusions but appeared more severely affected, so it was not possible to distinguish individual coxa, trochanter or femur segments (Fig 2D, and Fig. 3D,E). However, unlike the *Dll-Gal4*, some tarsal segments and the attached claws were present. We suspect that the differences in phenotype reflect differences in abundance, timing or position of expression of Gal4 drivers. Bristles on the *disco*-lof legs mostly lacked bracts (Fig. 3), suggesting they were of proximal leg type. In addition, leg bristle polarity was disrupted. The bristles were oriented somewhat randomly, with only small groups similarly aligned (Fig. 3). We confirmed these phenotypic results by removing the *disco* genes using clonal analysis (data not shown) with the same deficiency we used in our prior work (Patel et al., 2007), but this time targeting the center of the *disco*-expressing region using the *Dll-Gal4* driver to induce FRT recombination.

When a leg was completely lacking, the ventral body wall was also affected. For example, when a first thoracic leg was absent, the five small microchaetae along the ventral-anterior thorax were also absent (Fig 2B’-E’). When a second thoracic leg was absent, most sternopleural and mesosternal bristles were also absent (Fig. 2 C’-E’). In the most extreme case (Fig 2E) the fly had no legs and no discernable ventral body wall features except for a single bristle, likely a sternopleural.

**Figure 2:**
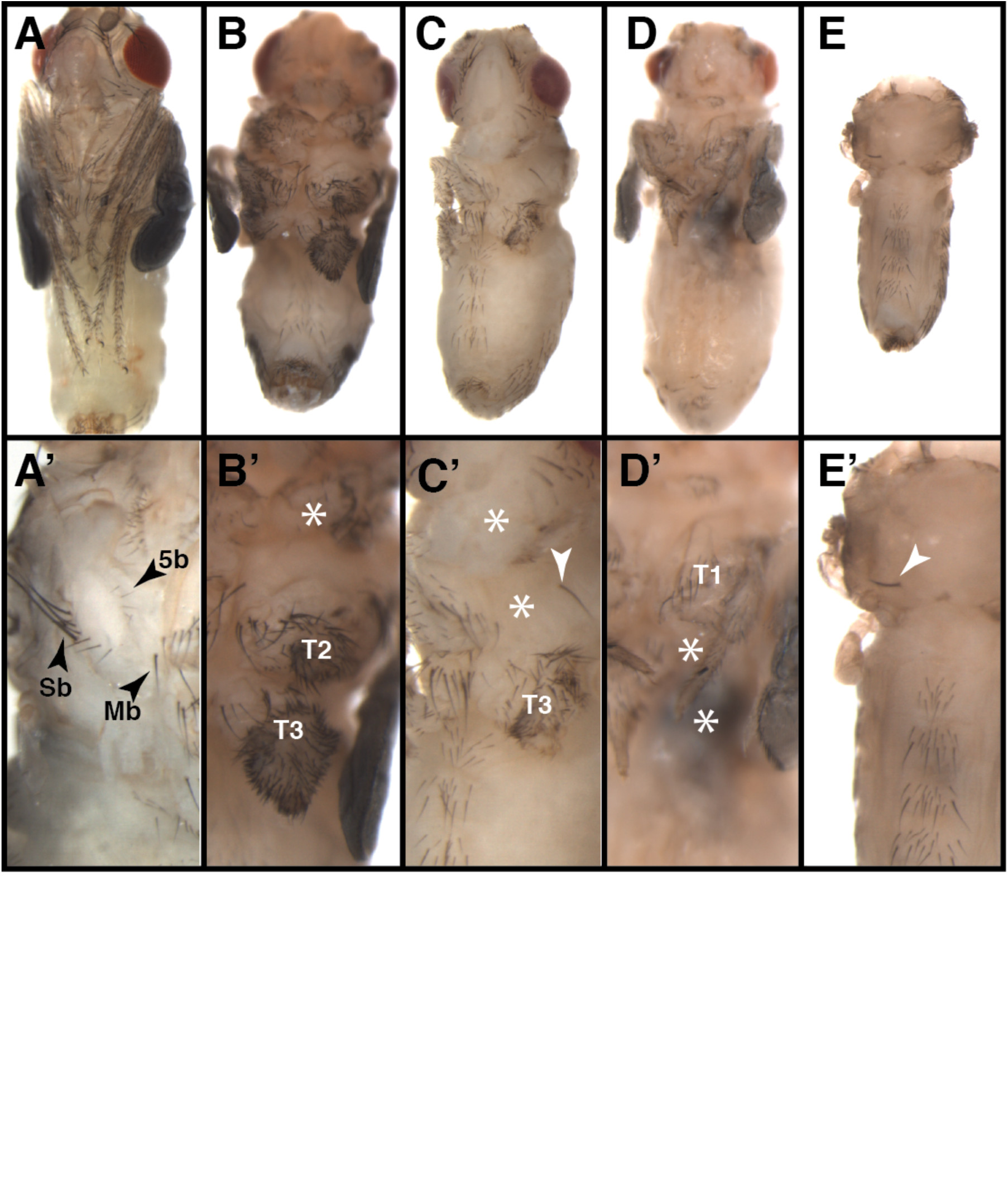
Phenotype of *disco*-lof pharate adults. (A-E) Ventral views of the flies, and (A’-E’) close-up views of the corresponding ventral thorax. (A, A’) Oregon R. In A’ the wings and legs have been removed to expose the characteristic bristles (arrowheads) of the ventral thorax: Sb, sternopleural bristles; Mb, mesosternal bristle; 5b, the five small microchaetae along the ventral-anterior in the first thoracic segment. (B-E and B’-E’) *disco*-lof flies (B) An example of *Dll-Gal4 disco*-lof pharate adult missing a first thoracic leg (asterisk in B’) with enlargement of the proximal portion of the second and third legs. (C-C’) *Dll-Gal4 disco*-lof pharate adult missing first and second thoracic legs (asterisks C’). A single bristle (arrowhead in C’) remained near to where the second leg should have been. (D-D’) *Act-Gal4 disco*-lof pharate adult. The first leg was reduced and the second and third legs were missing (asterisks in D’). Note that first thoracic leg maintained some tarsal structures including a few sex comb teeth. This was a hallmark of many *Act-Gal4 disco*-lof pharate adults. (E-E’) The most extremely affected fly lacked most ventral identity. Only a single bristle remained (arrowhead in E’).

**Figure 3:**
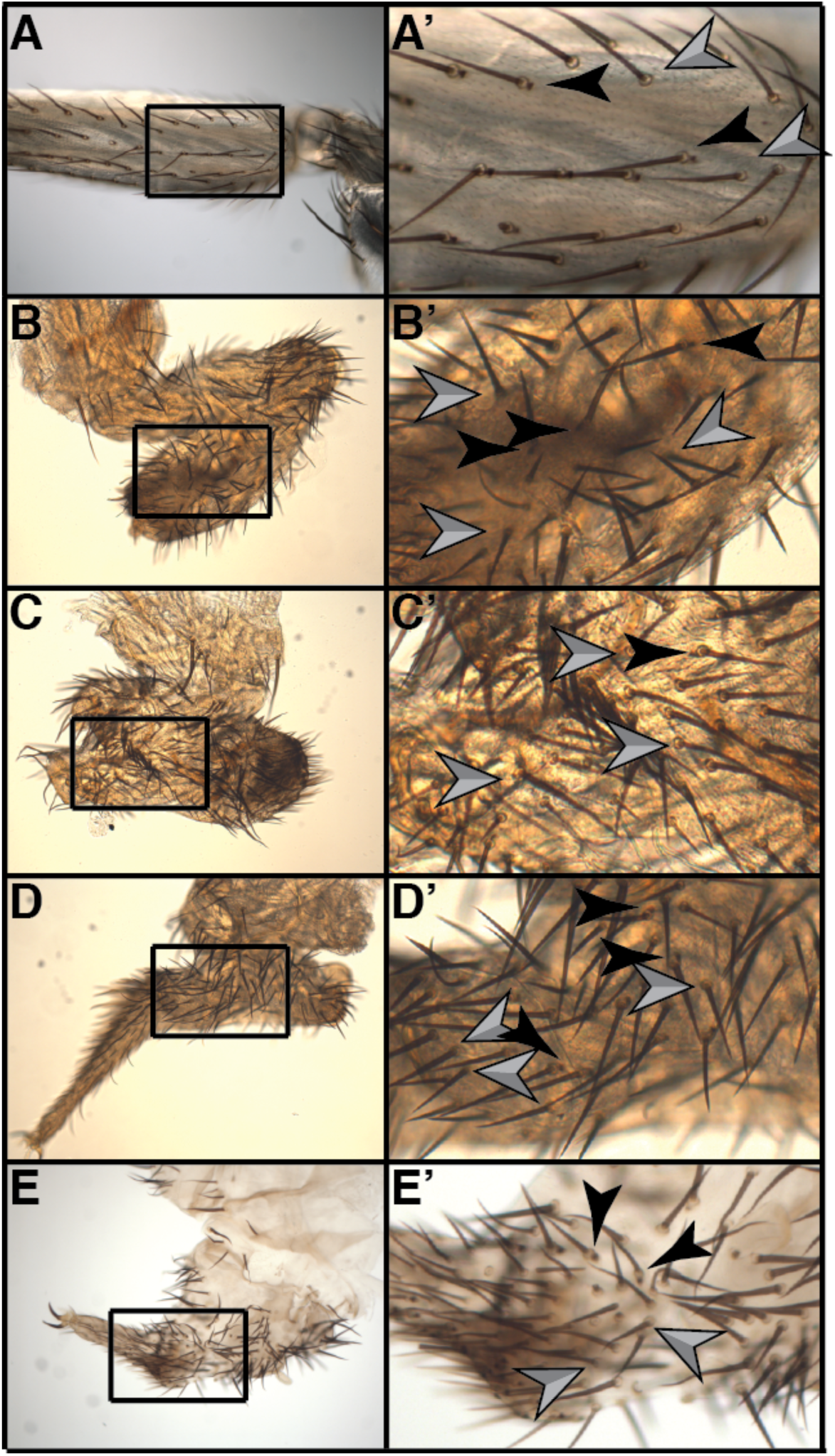
Phenotype of *disco*-lof pharate adult legs. Prime letters are close-ups of the corresponding legs. (A) Wild type femur with both bracted and non-bracted bristles. In general, non-bracted bristles are characteristics of the proximal portion of the legs and extend usually into the proximal femur as shown here. More distal to this, bracted bristles are most prevalent, distally in the femur, tibia and tarsi. (B-C) Legs from pharate adults of *Dll-Gal4* driving *disco*-lof and (D-E) *Act-Gal4* driving *disco*-lof. In all cases, bracted and non-bracted bristles were intermixed in the remnants of the affected legs, though non-bracted bristles were the predominant type. Gray arrowheads point to examples of bractless bristles, and black arrowheads point to examples of bracted bristles. Note that the orientation of the bristle was also disrupted.

### Morphology of *disco*-lof leg discs

Defects in the leg imaginal discs were apparent immediately upon inverting wandering third instar larvae. Though variable, the discs were often quite small when compared to those from their wild type siblings (Fig. 4). Wild type leg discs have a typical rounded, teardrop shape with a stalk dorsally (Fig. 4A). Characteristic folds, appearing as rings, demarcate future leg segments or groups of segments. In the center, the “end knob” will give rise to the most distal structures. The most extreme *disco*-lof discs were small, narrow structures with little obvious morphology (Fig. 4D,E). Less severely affected discs were reduced but retained some segmental folds (Fig. 4C). Some appeared only slightly smaller than wild type (Fig. 4B). The set of discs present in most larvae was a combination of both classes; the number of small versus reduced discs paralleled the number of missing versus truncated legs we observed in the pharate adults. Note, we did not consider rare cases where discs were completely missing, as discs could be torn off inadvertently during the dissections.

**Figure 4:**
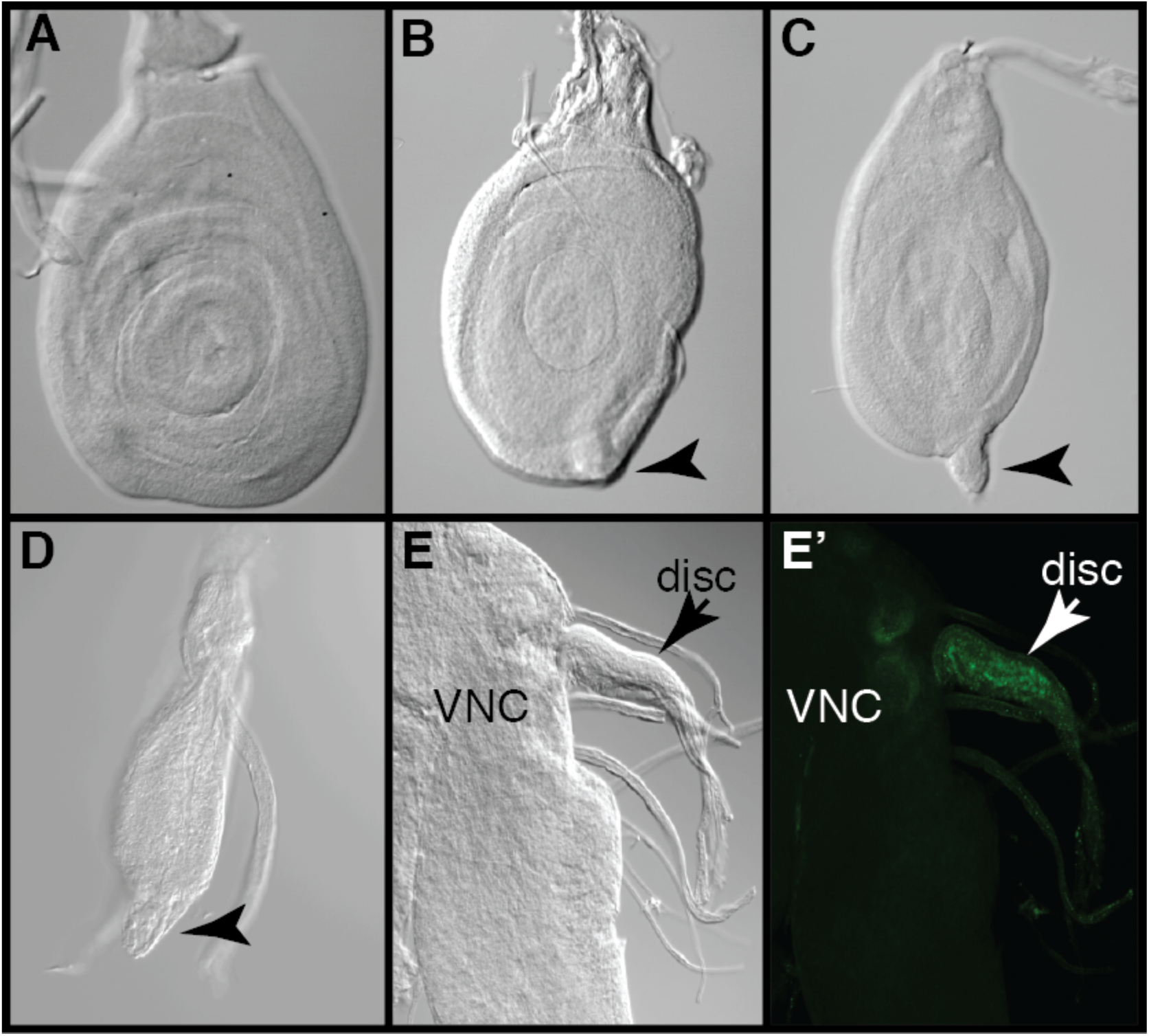
Morphology of *disco*-lof leg imaginal discs. (A) Leg disc of an Oregon R third instar larva. (B-E) A series of late third instar leg imaginal discs from *Act-Gal4*-driven *disco*-lof larvae. Note the variation in the sizes of the discs. (B) This leg disc was of a similar shape but somewhat reduced in size compared to that of the Oregon R (A). (C) This disc was about half the width of the Oregon R control and has a ventral protrusion, (arrowhead). (D and E) leg discs that were extremely reduced. In a severe case (E), the leg disc only reached a size slightly larger then the nerve entering the disc, which was still attached to the ventral nerve cord (VNC). (E’) Note that this extremely reduced disc still accumulated Tsh protein (green) though we did not detect other discs determinants that we stained for (data not shown).

### Gene expression in *disco*-lof leg discs

With the reduction in size of the legs and discs, we next examined the accumulation of several gene products that pattern the leg discs, Tsh protein (proximal) and Dac protein (medial) and *Dll* mRNA (distal) using Oregon-R as controls (see Materials and Methods).

In the most severely reduced discs we did not detect *Dll* mRNA or Dac protein (data not shown), though we did detect weak staining for the Tsh (Fig. 4E’). We noted that the Tsh-positive nuclei appeared more dispersed than those of the columnar cells in the wild type discs. In less severely affected discs, *Dll* mRNA was reduced in comparison to wild type (Fig. 5). Even in those discs that were nearly equal in size to wild type controls, *Dll* mRNA was quite reduced. Dac protein staining was occasionally reduced as well, particularly in smaller discs. We took intensity measurements from the images for all probes (Tsh, Dac and *Dll*) using three regions of brightest intensity within each disc (Fig. 5B). These measurements confirmed that *Dll* was significantly reduced (p=0.00004, Fig 5), and though Dac appeared to be reduced in some discs, the p value of those shown was just greater than the 5% probability level (p=0.06). There was no significant difference in Tsh staining between controls and these *disco*-lof discs.

**Figure 5:**
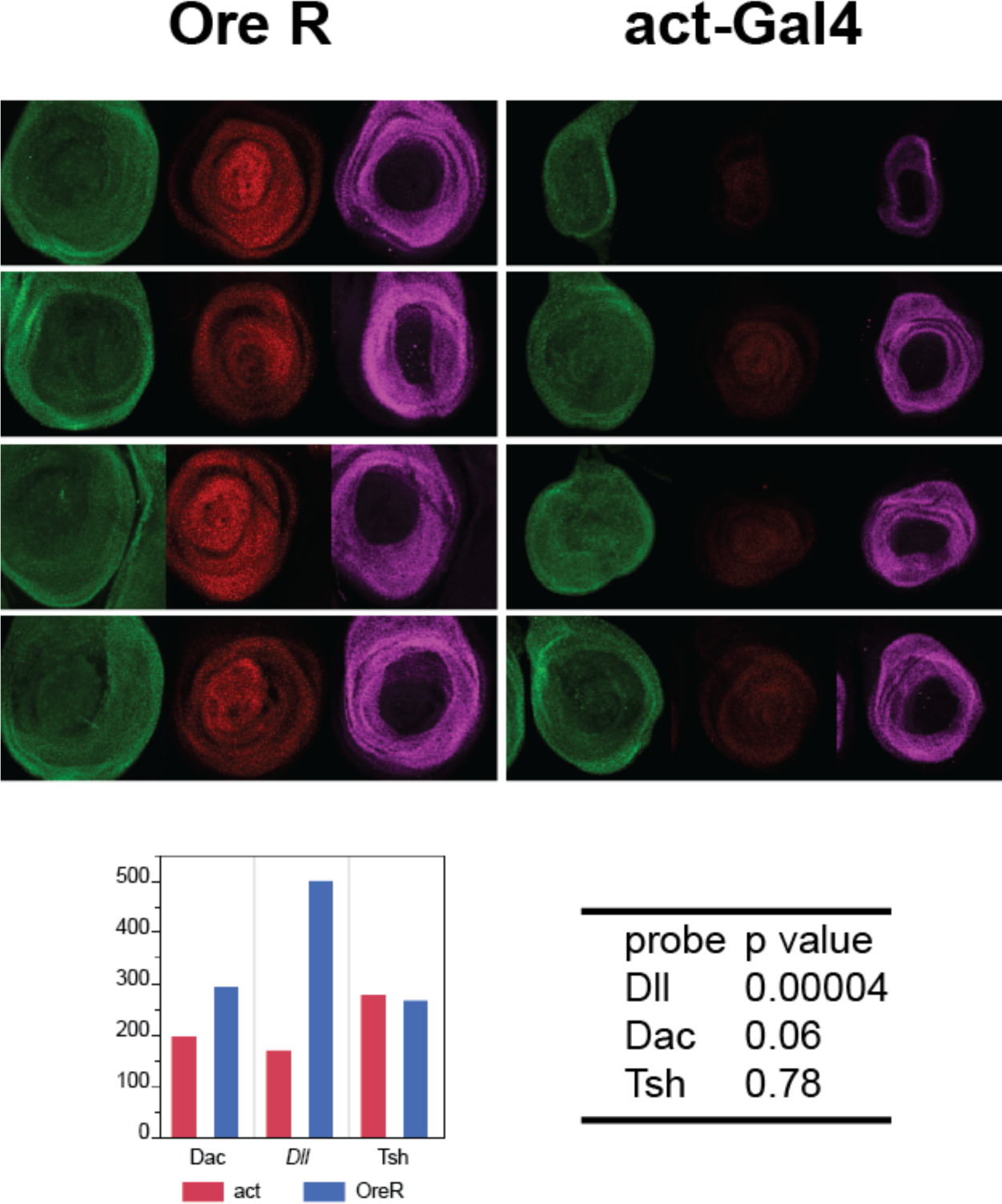
Analysis of leg gene determinants in *Act-Gal4 disco*-lof leg imaginal discs. We compared Tsh (green) and Dac (magenta) proteins and *Dll* transcript (red) accumulations between Oregon R and the *disco*-lof genotype using ImageJ (Schneider et al., 2012) to obtain estimations of fluorescence abundance (see materials and methods). The graph is in arbitrary units. Tsh measurements were nearly identical between the two genotypes, while Dac was reduced, but was just above significance cutoff of 0.05. On the other hand, the difference in *Dll* RNA accumulation was statistically significant (P-value = 0.00004) when compared to Oregon R.

*Dll* (Lecuit and Cohen, 1997) and, indirectly, *dac* (Giorgianni and Mann, 2011) are regulated by Wg and Dpp in the leg discs, and since reduction of either Wg or Dpp can cause truncation of the distal legs, it seemed plausible that expression or function of these morphogens could be altered in these discs. Therefore, we assayed *wg* transcript and protein and *dpp* transcript and activity (via p-mad) in *disco*-lof discs of similar size to those that had reduced *Dll* mRNA. In control discs *wg* mRNA and protein accumulate in a wedge-shape sector in the anterior-ventral portion of the leg disc, extending along the disc folds into the posterior compartment (Fig. 6A-A’’’). Wg protein was more widely dispersed than the mRNA (Fig. 6A’’’). In moderately affected *disco*-lof discs, *wg* mRNA did not extending as far anteriorly around the disc and had a sharper boundary along the posterior edge (Fig. 6B-B’). Wg protein (Fig. 6B’’) was reduced overall and did not appear to spread far from the area of transcript accumulation (Fig. 6B’’’). By contrast, *Dpp* transcript and function appeared to be unaffected (data not shown). However, in the most severely affected discs, those that were highly reduced in size, we did not detect either *wg* (6C-C’’’) or *dpp* (data not shown).

**Figure 6:**
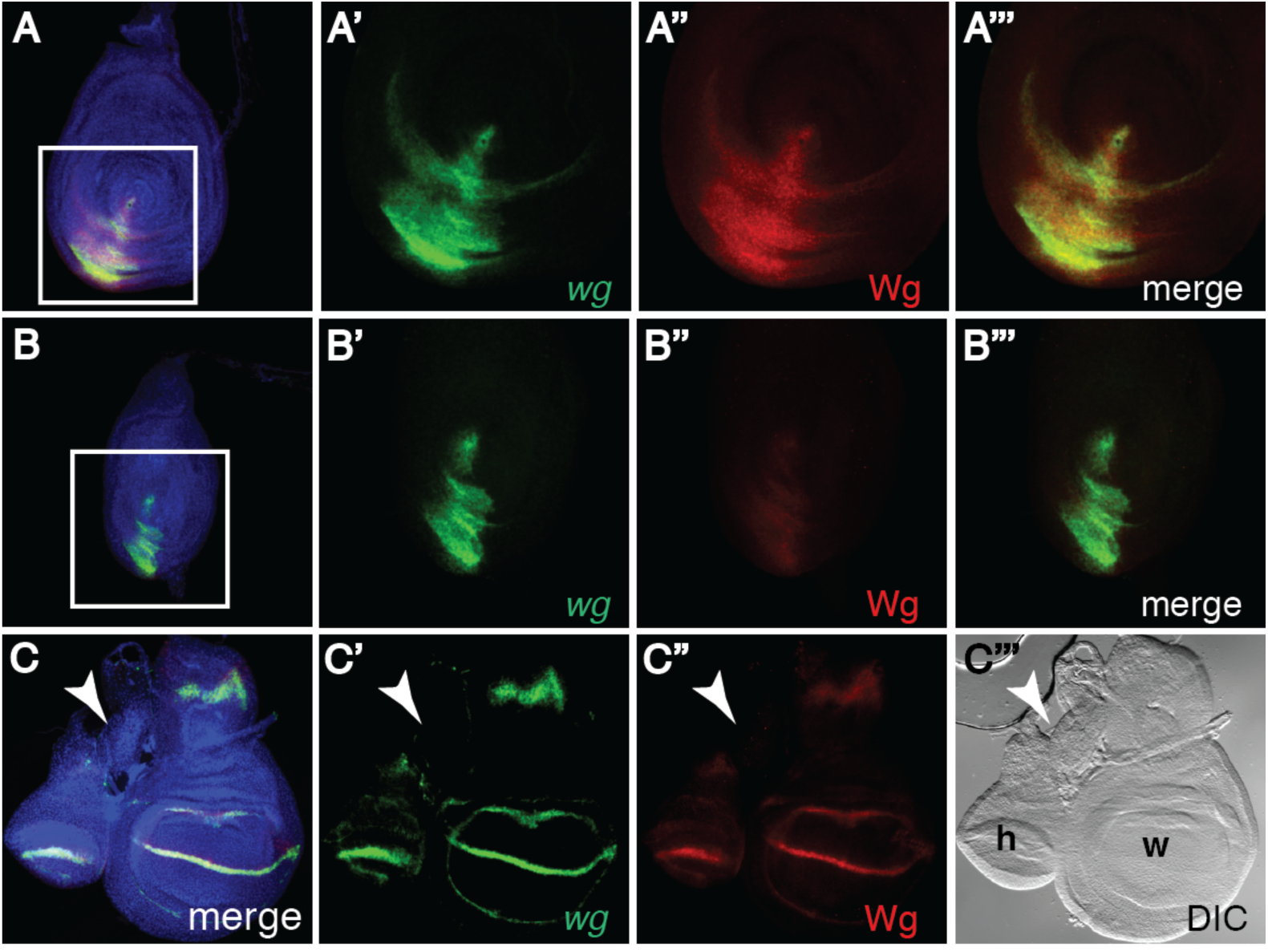
Analysis of *wingless* transcript and protein accumulation in *disco*-lof leg imaginal discs. (A-A’’’) Oregon R leg disc stained to detect wg transcripts (green) and Wg protein (red). A’-A’’’ are enlargements of the marked area in A. (B-C’’’) *Act-Gal4 disco*-lof discs. B’-B’’’ are enlargements of the region marked in B. In *disco*-lof discs *wg* transcripts appear to be more restricted (B’) when compared to Oregon R (A’). The accumulation of Wg protein is likewise restricted, but also appears to be reduced (B’’, B’’’) when compares to Oregon R (A’’, A’’’). Note that in Oregon R, Wg protein spreads further than the area of transcript (A’’’), but this is not the case in the *disco*-lof discs (B’’’). A more severely affected *disco*-lof third thoracic leg disc is shown in C-C’’’(arrowhead). We did not detect *wg* RNA or protein in discs such as this, though both appeared normally in the wing and haltere. The DIC image shows the size relationship between the haltere (h), the wing discs (w) and the leg (white arrowhead) discs from the same third instar larva. Discs in A and B are oriented dorsal up and anterior to the left, while C have the posterior to the left. All magnifications were 20X except for C-C’’’ which were 10X so as to include the haltere and wing for comparison.

### Cell Death in *disco*-lof discs

In our prior report, we proposed that leg disc cells lacking the *disco* genes may not be viable (Patel et al., 2007). Deletion of leg segments or entire legs and reduced size of the *disco*-lof leg discs could also indicate that cells of the affected discs have become weakened and may be undergoing apoptosis. To determine if this were the case, we performed TUNEL staining. In *disco*-lof discs, we observed an increase in TUNEL positive cells compared to wild type (Fig 7A). We noted increased TUNEL staining in *disco*-lof discs of near normal size and in those severely reduced (Fig 7B,C, respectively). TUNEL stained cells were scattered somewhat randomly throughout the discs. Many were located in the lumen of the disc, suggesting they were extruded from the discs, as is common of apoptotic cells (Rosenblatt et al., 2001). Only an occasional TUNEL positive cell was detected in wild type discs (Fig. 7A).

**Figure 7:**
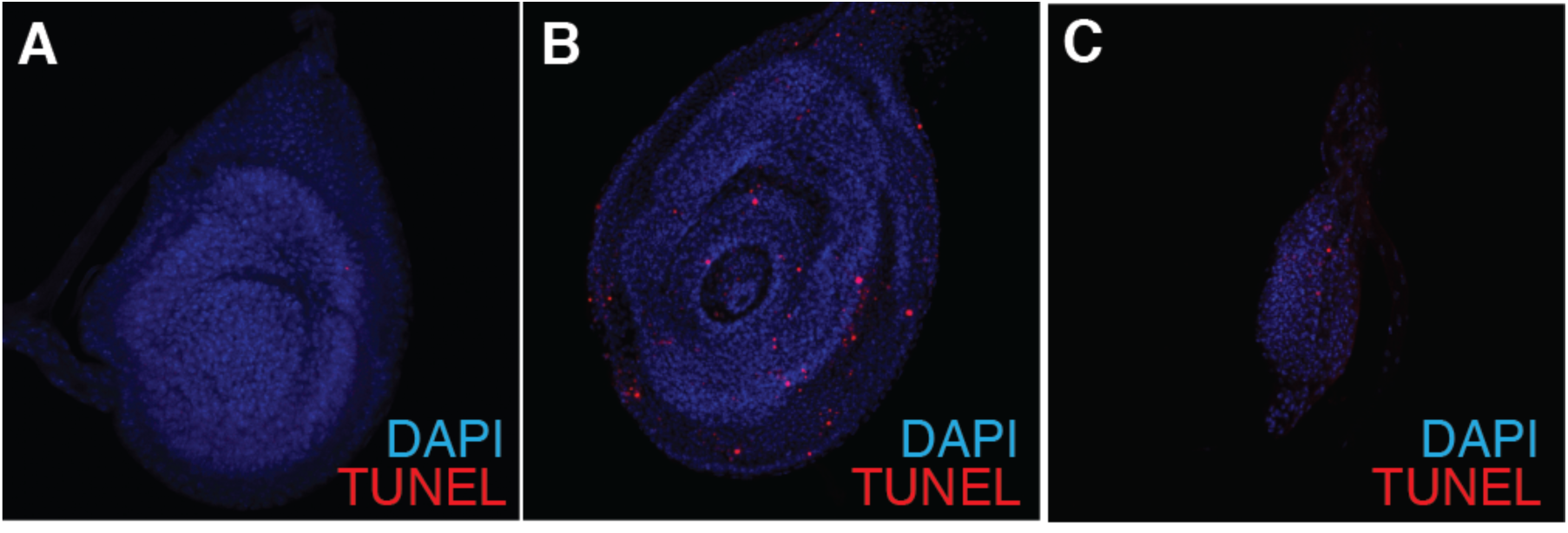
TUNEL analysis of *disco*-lof leg discs. (A) Leg imaginal disc of an Oregon R fly. We detected one TUNEL-positive cell. (B) A nearly normal sized *disco*-lof leg disc. These discs had an increase in TUNEL staining, as did more severely affected leg discs (C). In both B and C many of the TUNEL positive cells were detected in the lumen of the discs.

### Establishment of the leg disc primordia during embryogenesis

Cells of the leg discs primordia are set aside during early embryogenesis; their positions marked by the expression of *Dll* (Cohen et al., 1991). A previous study (Cohen et al., 1991) compared *disco* and *Dll* expression in the emerging primordia and concluded that *disco* was expressed after *Dll*, and, further, that in the absence of *Dll*, *disco* was not activated. However, this was based on expression of the enhancer trap line C50.1S1 to represent *disco* expression, and we have previously shown this enhancer trap to be inserted into the *disco-r* gene, not *disco* (Patel et al., 2007). Therefore, we reexamined *Dll* and *disco* expression in early embryos. The thoracic region of wild type embryos at two different stages of development stained to detect *disco* and *Dll* mRNAs are shown in Fig 8. Both genes appear to accumulate at stage 10/11 (Fig. 8 A-A’’’), where their transcripts overlapped in the primordia. We could not detect a difference in their distribution. However, *Dll* expression is dynamic (Cohen et al., 1993; Estella et al., 2012; Galindo et al., 2011; McKay et al., 2009; O’Hara et al., 1993; Vachon et al., 1992), and, though at this early stage *Dll* accumulates throughout the thoracic appendage primordia, by late stage 12, *Dll* transcripts are restricted to what will be the medial to distal portion of the legs. At this stage, *disco* mRNA was more broadly distributed (Figure 8 B-B’’’), and this remained the case throughout the rest of development, including in the third instar leg discs (Patel et al., 2007 and data not shown).

**Figure 8:**
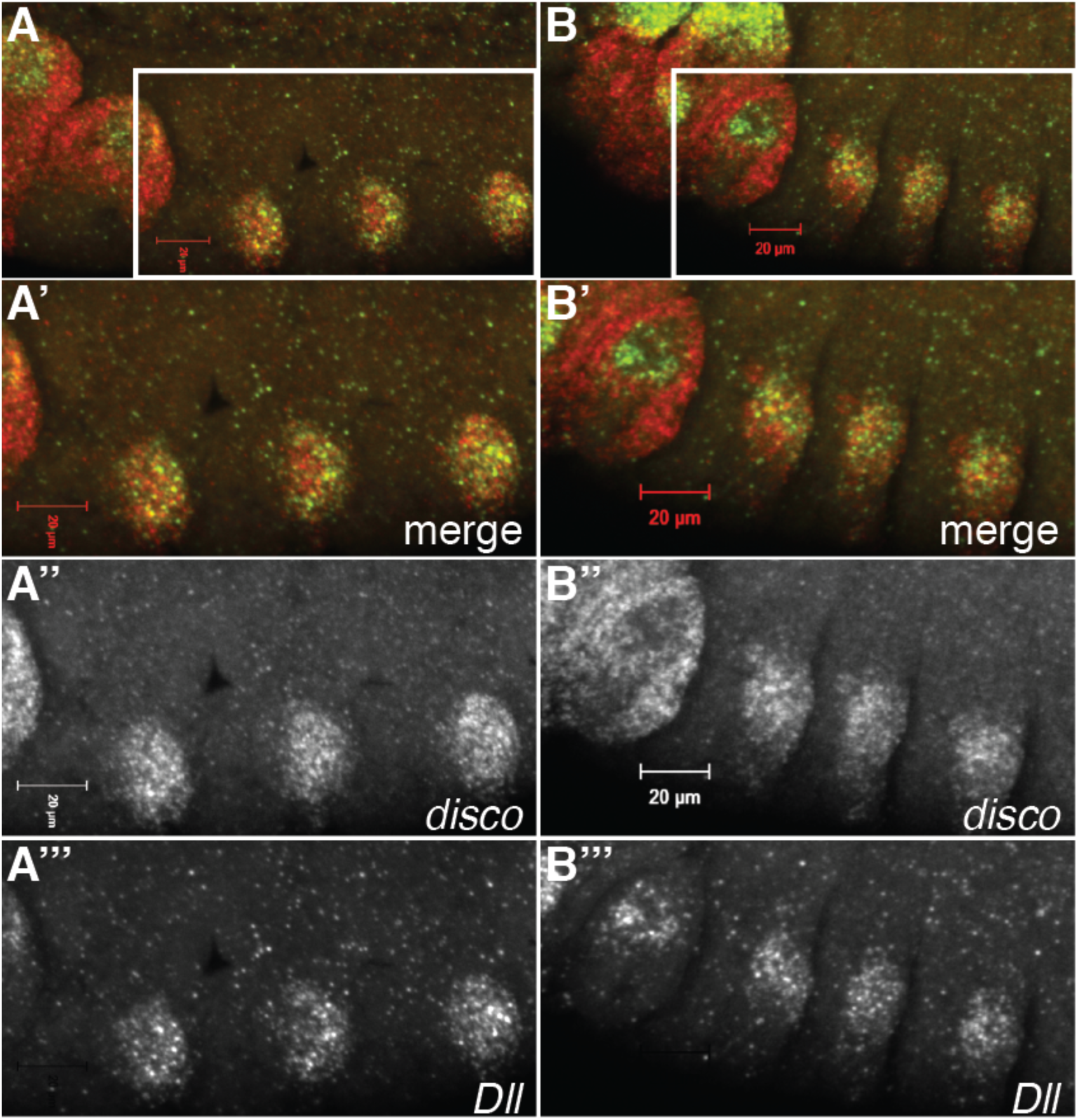
*disco* and *Dll* transcript localizations in wild type Oregon R embryos at two different developmental stages. (A-A’’’) Embryos at stage 10/11, (B-B’’’) embryos at late stage 12. *disco* RNA is in red and *Dll* RNA in green. Prime letters indicate enlargements of the marked rectangles in A and B. Scale bar at 20µm. During early embryogenesis (A-A’’’) *disco* (A’, A’’) and *Dll* (A’, A’’’) transcripts were detected in the same groups of cells in the imaginal disc primordia. However, at the later stage (B-B’’’) *disco* transcripts (B’, B’’) were detected more broadly than *Dll* (B’, B’’’), particularly noted toward the ventral portion of the leg disc primordia. Dorsal is up and anterior to the left.

Cis-regulatory modules have been identified at the *Dll* gene that mimic *Dll* regulation (Cohen et al., 1993; Estella et al., 2008; Estella et al., 2012; Galindo et al., 2011; McKay et al., 2009; O’Hara et al., 1993; Vachon et al., 1992). For example, the *304* element is initiated by Wg during stage 10-11 of embryogenesis throughout the thoracic appendage primordia. This enhancer is only active for a brief period. Later, at stage 12-13, enhancers such as *LP* and *LT* take over. Unlike 304, these enhancers only drive expression in the medial to distal regions of the leg discs (Galindo et al., 2011; McKay et al., 2009) that will become the *Dll* and *dac* domains of later leg discs (Estella et al., 2008; Estella et al., 2012).

Given the timing of *LT* activation and its dependence upon both Wg and Dpp signaling (Estella et al., 2008) we questioned whether *LT* would react to the loss of the *disco* gene functions. To test this, we examined ß-galactosidase production by the *LT* reporter in embryos hemizygous for *Df(1)ED7355* which deletes both *disco* and *disco-r*. At stage 16 in wild type embryos, *LT* is expressed in a ring of cells in the leg disc primordia (Fig.9 A and see McKay et al., 2009). In embryos hemizygous for *Df(1)ED7355*, LT expression was weaker and in fewer cells (Fig. 9B). The normal ring-shaped pattern was disrupted. We noted no change in the thoracic expression of the *304* reporter in Df(1)ED7355 embryos, though interestingly, it was derepressed in some head segments (data not shown).

**Figure 9:**
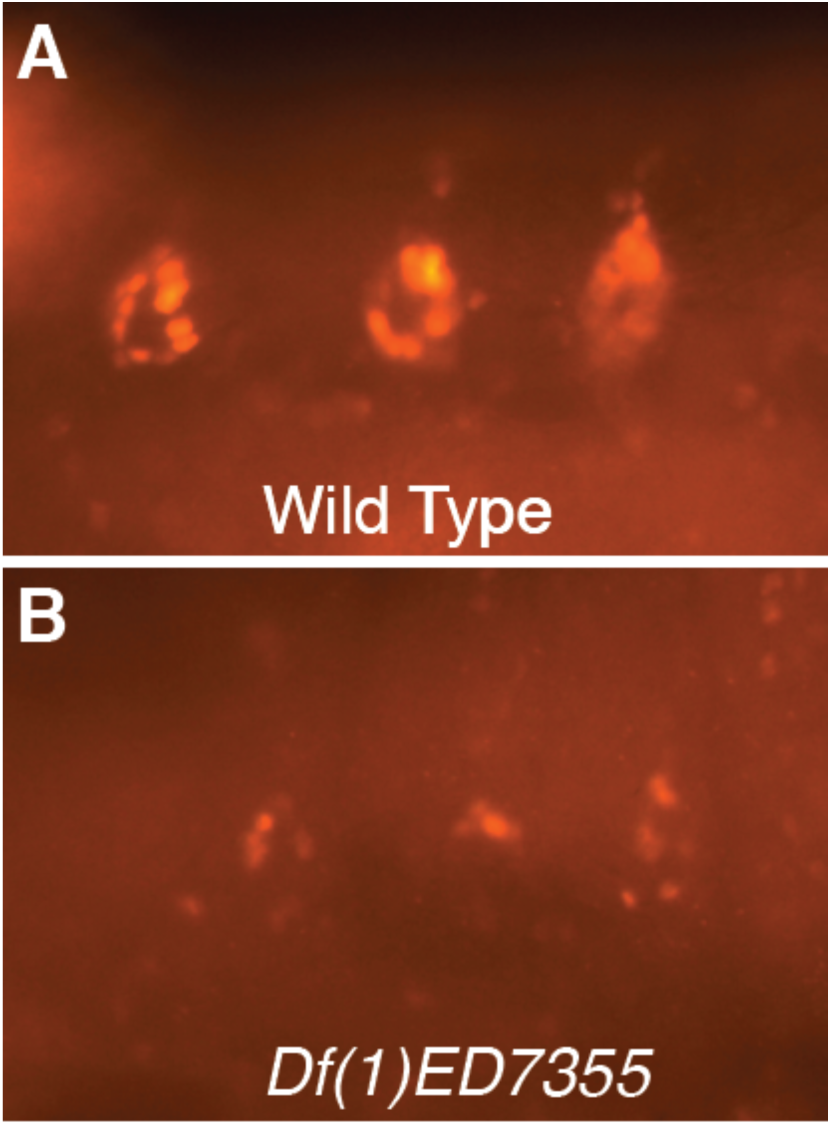
*LT* driven Beta-gal accumulation in wild type and *Df(1)ED7355* hemizygous embryos lacking *disco* and *disco-r*. In wild type embryos (A) *LT* drives Beta-gal accumulation in rings of cells in the thoracic segments marking cells of the leg imaginal disc primordia. In *Df(1)ED7355* hemizygous male embryos (*Df(1)ED7355*/*Y*), only a few cells stained in each hemisegment, and they were not organized in rings. It is unknown whether these cells are true disc primordia cells or perhaps other cells in the region that activated *LT*. The identity of the *Df(1)ED7355* hemizygous embryos was confirmed by simultaneous detecting *disco* mRNA in the embryo collection, which was absent in the hemizygous embryos due to the deficiency. Dorsal is up and anterior to the left.

## DISCUSSION

Appendage development in Drosophila has been extensively studied as a model system to decipher the genetic control of pattern formation during animal development. Previous studies have led to a model whereby signaling systems in the embryo establish the leg primordia, followed by patterned gene expression that establishes the overall axes of the leg and subsequently specifies leg differentiation (see Fig. 1). Previously, we demonstrated that the genes *disco* and *disco-r* appear to be major factors in leg development, particularly since ectopic expression of either gene in the wing imaginal disc could induce legs in place of wings (Patel et al., 2007). However, until now we were unable to ascertain the requirement of these genes during development of the adult leg.

Removing the *disco* gene functions from the second instar onward led to two defects. In the most severe cases, all but the most proximal body wall tissues was missing. Many discs were reduced to thin slivers of tissue having little, if any, accumulation of canonical leg determinants, with the exception that a few cells accumulated Tsh. Since the body wall arises from the peripodial cells and cells from the marginal zone of the discs (the region between the columnar disc cells and peripodial epithelial cells, Fristrom and Fristrom, 1993; McClure and Schubiger, 2005; Milner et al., 1984), and since cells in this marginal zone express the *disco* genes (Patel et al., 2007, and data not shown), we suspect that these highly reduced discs were incapable of producing any overt appendages, though they may be able to contribute to the body wall of the ventral thorax. Considering our observations, we suggest that the *disco* genes function as positional determinants, much like *tsh*, *Dll* and *dac*, but having a larger domain that encompasses all but the most proximal portion of the leg disc derivatives.

Our observations of less severely affected legs and discs were just as informative. Examining these legs and discs indicated that the Disco transcription factors might be necessary to stabilize/maintain the expression of other leg patterning genes, including factors for signaling and transcription, since expression of several genes were not well maintained once the Disco transcription factors were reduced.

It is not clear why the two phenotypes, leg truncations and complete loss, would be present in the same pharate adult. Perhaps this indicates that there were slight differences in efficacy of the *disco-r* RNAi in different leg discs. If so, this could indicate that level of the residual Disco transcription factors was responsible. However, it could also indicate that there is a change in the requirement of these proteins at about this stage, and indeed, there is a change in gene regulation at this time.

From these studies, we propose that the *disco* gene products are particularly necessary during transitions in gene expression during leg determination and development. Initiation of the thoracic disc primordia begins at embryonic stage 11, when the segment polarity signal Wg induces small groups of cells in each thoracic hemisegment to express appendage specification genes, such as *Dll* and the *disco* genes among others (Cohen et al., 1993; Cohen, 1990; Kubota et al., 2003). It is important to note that at this early stage these groups of cells include the primordia of the ventral (leg) and dorsal (humeral, wings and halters) thoracic discs, and also the larval Keilin’s Organs (vestiges of larval legs) (Angelini and Kaufman, 2005; Bolinger and Boekhoff-Falk, 2005; Cohen et al., 1991; McKay et al., 2009; Panganiban, 2000b). Loss of *disco* and *disco-r* does not appear to alter the initial induction of the thoracic disc primordia. Indeed, the *Dll* and *disco* genes appear to be activated simultaneously, so it would be unlikely that one would be required for induction of the other. Furthermore, loss of the *disco* genes did not significantly alter expression of the *Dll-304* element, which responds to Wg induction (data not shown).

The first transition occurs at about embryonic stage 14 when the primordia of the ventral and dorsal discs (and the Keilin’s organs) separate so that each becomes an independent unit (Bolinger and Boekhoff-Falk, 2005; Cohen et al., 1991; McKay et al., 2009; Panganiban, 2000a). At this time, *Dll* is repressed in the dorsal thoracic discs and is restricted to only those cells that will form the telopodite region of the leg (the trochanter and more distal leg segments) (McKay et al., 2009). Significantly, Dpp signaling is now part of the activation process, where it was repressive during initiation (Campbell et al., 1993; Diaz-Benjumea et al., 1994; Goto and Hayashi, 1997; Kubota et al., 2000; Lecuit and Cohen, 1997). Cells in the leg discs that continue to express *Dll* must receive high levels of both Wg and Dpp signals (see Figure 1 above). This transition in gene regulation is reflected by changes in enhancer function. For example, the *Dll-304* enhancer, which responds to the Wg signaling during induction, is no longer active, and the *LT* enhancer, in part, directs *Dll* expression in telopodite region of the leg (Cohen et al., 1993; Estella et al., 2008; Galindo et al., 2011; McKay et al., 2009). Furthermore, the DKO enhancer takes over expression of *Dll* in the Keilin’s organs (McKay et al., 2009). Our observation that, the *LT* enhancer is not stably expressed in the absence of the *disco* gene products, indicating that the Disco transcription factors might be necessary to stabilize gene expression orchestrated by *LT*.

A second transition in gene regulation occurs during the late second instar larval stage, when expression of genes such as *Dll* become independent of Wg and Dpp (72-84 hours after egg laying, Estella et al., 2012; Galindo et al., 2002). This transition is also seen in *Dll* regulatory modules as the *LT* enhancer now requires the *M* element to remain active (Estella et al., 2008; Galindo et al., 2011, reviewed in (Estella et al., 2012). It is around this stage that flies with reduced *disco* gene products were able to survive long enough to produce the pharate adults, albeit with severe leg defects.

Whether the changes in gene expression in the *disco*-lof leg discs reflect direct regulation of genes by the Disco transcription factors will need to be addressed in the future. Interestingly, in the study by Negre et al. (2011), Disco binding sites were found near *Dll*, *dac*, and *wg*, and though no Disco binding sites were reported in the *Dll-LT* enhancer, we have discovered sequences in this element that closely resemble those bound by the Disco protein (Rosario and Mahaffey, unpublished). Further work will be necessary to determine whether these genes are direct targets of Disco and Disco-r.

Our loss of function analyses continue to show that the Disco proteins are major factors of the Drosophila leg and ventral appendage development pathway. These genes are members of a conserved family of genes encoding zinc finger transcription factors with homologs found in many animals (Knight and Shimeld, 2001). Considering the structural conservation, it is reasonable to ask whether the homologs have similar developmental roles. Indeed, our earlier work established that the single Tribolium homologue, *Tc-disco*, specified portions of the legs and other ventral appendages in this beetle (Patel et al., 2007). Further support for a conserved role comes from a report uncovering a possible link between a break in one of the human *disco* homologs, Basonuclin-2, and limb growth in humans (Bhoj et al., 2011). If this is indeed a causative mutation, then it is possible that the appendage role of the *disco* genes is conserved. Perhaps it is noteworthy as well that, given the role of Hh and *disco* in patterning Drosophila legs, Basonuclin is activated in some basal cell carcinomas by Gli proteins, the transcriptional responders of the Hh pathway (Cui et al., 2004). Indeed, future work will undoubtedly look to determine if there are such conserved roles and regulations.

## ACKNOWLEDGEMENTS

We thank the TRiP at Harvard Medical School (NIH/NIGMS R01-GM084947) for providing transgenic RNAi fly stocks in this study. Stocks obtained from the Bloomington Drosophila Stock Center (NIH P40OD018537) were used in this study. We also thank the Vienna Drosophila Resource Center (Dietzl et al., 2007) for providing fly lines. We thank Dr. Patricia Estes for the GFP fly line and Dr. Stephen Cohen for the Tsh antibodies. The Dachshund monoclonal antibody, mAbdac2-3 deposited by G. M. Rubin was obtained from the Developmental Studies Hybridoma Bank, created by the NICHD of the NIH and maintained at The University of Iowa, Department of Biology, Iowa City, IA 52242. This work was supported by a grant (IOS-0842201) from the Developmental Mechanisms Program of the National Science Foundation (www.nsf.gov) to JWM. No competing interests declared.

## AUTHOR CONTRIBUTIONS

JBR and JWM both developed the ideas, designed experiments and interpreted the data, and co-wrote the manuscript.

